# Cation-chloride cotransporters, Na/K pump, and channels in cell water and ions regulation: in silico and experimental studies of the U937 cells under stopping the pump and during RVD

**DOI:** 10.1101/2021.07.10.451878

**Authors:** Valentina E. Yurinskaya, Alexey A. Vereninov

## Abstract

Cation-coupled chloride cotransporters play a key role in generating the Cl‾ electrochemical gradient on the cell membrane which is important for regulation of many cellular processes. However, the cooperation of transporters and channels of the plasma membrane in holding the ionic homeostasis of the whole cell remains poorly characterized because of the lack of a suitable tool for its computation. Our software successfully predicted in real-time changes in the ion homeostasis of U937 cells after stopping the Na/K pump, but so far considered the model with only NC cotransporter. Here the model with all main types of cotransporters is used in computation of the rearrangements of ionic homeostasis due to stopping the pump and associated with the regulatory volume decrease (RVD) of cells swollen in hypoosmolar medium. The parameters obtained for the real U937 cells are used. Successful prediction of changes in ion homeostasis in real-time after stopping the pump using the model with all major cotransporters indicates that the model is reliable. Using this model for analysis RVD showed that there is a “physical” RVD, associated with the time-dependent changes in electrochemical ion gradients, but not with alteration of channels and transporters of the plasma membrane that should be considered in studies of truly active regulatory processes mediated by the intracellular signaling network. The developed software can be useful for calculation of the balance of the partial unidirectional fluxes of monovalent ions across the cell membrane of various cells under various conditions.

## INTRODUCTION

The role of Cl‾ channels and transporters in cellular processes attracts much attention at present (Hoffmann et al., 2015); Jentsch 2016; Pedersen et al., 2016; Jentsch, and Pusch, 2018; Currin et al., 2020; Murillo-de-Ozores et al., 2020). Cation-coupled chloride cotransporters of the SLC12 family play a key role in generating the Cl‾electrochemical potential difference on the cell membrane which is mandatory for Cl^-^‾ and Cl‾ channels to be a regulator of cell volume, intracellular pH and cell signaling (Gamba, 2005, 2009). The progress in molecular studying cation-coupled chloride cotransporters and Cl‾ channels is impressive. However, the current studies in this area focus mostly on the regulation of channels and transporters but not an analysis of their interactions in maintaining the entire ionic homeostasis of cell, regulation of the cell water balance and generation of electrochemical gradients of ions on the cell membrane (Hoffmann, Pedersen, 2010; Cruz-Rangel et al., 2012; Kaila et al., 2014; Zhang et al. 2016; Shekarabi et al., 2017; de Los Heros et al., 2018; Wilson, and Mongin, 2018; Dmitriev et al., 2019; Okada et al., 2019; Song et al., 2019; Bortner and Cidlowski, 2020; Kittl et al., 2020; Murillo-de-Ozores et al., 2020; Pacheco-Alvarez et al, 2020). We believe that this is partly due to the lack of a suitable computational modeling tool for a rather complex system. The software for calculating the balance of unidirectional fluxes of monovalent ions via main ion pathways in the cell membrane has been developed by us in recent years (Vereninov et al., 2014; 2016; Yurinskaya et al., 2019). The software was supplied by a simple executable file that allowed, based on the minimum necessary experimental data, to find all the characteristics of ion homeostasis and a list of all unidirectional fluxes of monovalent ions through the main pathways in the cell membrane. Until now we tested our tool in prediction of rearrangement of ion homeostasis in U937 cells caused by stopping the Na/K pump using the incomplete model with only NC cotransporter Vereninov et al., 2014; Yurinskaya et al., 2019). The model with all major types of cotransporters for apoptotic U937 cells was considered in our recent study (Yurinskaya et al., 2020) which showed that the effects of KC and NKCC are small. The first goal of the present study was to find out whether a model with a full set of cotransporters would be successful in predictions of changes in ion homeostasis after stopping the pump. The data obtained allow us to conclude that the tool and model are reliable. It was also interesting to apply the same approach to the analysis of changes in the ionic homeostasis of cells placed in a hypoosmolar medium, when, as a rule, a regulatory volume decrease (RVD) occurs after rapid swelling. The RVD phenomenon has attracted a lot of attention for more than half a century (Hoffmann et al., 2009; Hoffman, Pedersen, 2010). However, no mathematical modeling of RVD, considering all the main cotransporters and based on the real parameters of the cells, has yet been carried out. Study of RVD in the U937 cell model presented below reveals many interesting effects in changing the unidirectional fluxes of Na^+^, K^+^, Cl‾and in the whole ionic homeostasis during RVD that occur without any changes in properties of channels and transporters of the cell membrane. The effects that can be qualified as a “physical” RVD mask truly active regulatory processes mediated by the intracellular signaling network, cell kinases etc. Using our software allows to separate the effects of changing external osmolarity, cell water balance and electrochemical potential differences driving ions across the cell membrane from the effects caused by changes in properties of the membrane channels and transporters. The modeling helps to quantify the effects caused by alteration of each ion pathway separately and in combination more rigorously than using specific inhibitors or mutation analysis.

## MATERIALS AND METHODS

### Cell cultures and solutions

Human histiocytic lymphoma U937 cells, myeloid leukemia K562 cells, and T lymphocyte Jurkat cells were obtained from the Russian Collection of Cell Cultures (Institute of Cytology, Russian Academy of Sciences). The cells were cultured in RPMI 1640 medium supplemented with 10% FBS at 37 °C and 5% CO_2_ and subcultured every 2-3 days. Cells, with a culture density of approximately 1 × 10^6^ cells per ml, were treated with 10 µM ouabain or hypotonic 160 mOsm solution. A stock solution of 1 mM ouabain in PBS was used. A hypoosmotic solution was prepared from an isoosmotic medium by decreasing the NaCl concentration by 75 mM, namely by mixing a standard isotonic medium with a medium of the same composition only without NaCl. Isoosmolar medium replacing 75 mM NaCl with 150 mM sucrose was prepared using a stock solution of 2 M sucrose in PBS. The osmolarity of all solutions was verified with the Micro-osmometer Model 3320 (Advanced Instruments, USA). All the incubations were done at 37 °C.

### Reagents

RPMI 1640 medium and fetal bovine serum (FBS, HyClone Standard) were purchased from Biolot (Russia). Ouabain was from Sigma-Aldrich (Germany), Percoll was purchased from Pharmacia (Sweden). The isotope ^36^Cl‾ was from “Isotope” (Russia). Salts and sucrose were of analytical grade and were from Reachem (Russia).

### Cellular ion and water content determination

The analysis of intracellular ion and water content is described in detail in our previous studies (Yurinskaya et al., 2005, 2011; Vereninov et al., 2007, 2008). Briefly, intracellular K^+^, Na^+^, and Rb^+^ were determined by flame emission using a Perkin-Elmer AA 306 spectrophotometer, intracellular Cl‾ was determined using a ^36^Cl‾ radiotracer. Cell water content was estimated by the buoyant density of the cells in continuous Percoll gradient, as *v*_prot._= (1-*ρ/ρ*_dry_)/[0.72(*ρ*-1)], where *ρ* is the measured buoyant density of the cells and *ρ*_dry_ is the cell dry mass density, which was given as 1.38 g ml^-1^. The share of protein in dry mass was given as 72%. Note that the relative changes in the water content in cells do not depend on the accepted values of the density of the dry mass of cells and the proportion of protein in it. The content of ions in the cell was calculated in micromoles per gram of protein, and the content of water in ml per gram of protein.

### Statistical analysis

Data are presented as the mean ± SEM. P < 0.05 (Student’s t test) was considered statistically significant.

### The mathematical background of the modeling

The mathematical model of the movement of monovalent ions across the cell membrane was similar to that used by Jakobsson (1980), and Lew with colleagues (Lew, Bookchin, 1986; Lew et al. 1991), as well as in our previous works (Vereninov et al., 2014, 2016; Yurinskaya et al., 2019, 2020). It accounts for the Na/K pump, electroconductive channels, cotransporters NC, KC, and NKCC. In this approach, the entire set of ion transport systems is replaced by a reduced number of ion pathways, determined thermodynamically, but not by their molecular structure.

All the major pathways are subdivided into five subtypes by ion-driving force: ion channels, where the driving force is the transmembrane electrochemical potential difference for a single ion species; NKCC, NC and KC cotransporters, where the driving force is the sum of the electrochemical potential differences for all partners; and the Na/K ATPase pump, where ion movement against electrochemical gradient is energized by ATP hydrolysis. This makes it possible to characterize the intrinsic properties of each pathway using a single rate coefficient. The following abbreviations are used to designate ion transporters: NKCC indicates the known cotransporters of the SLC12 family carrying monovalent ions with stoichiometry 1Na^+^:1K^+^:2Cl‾ and KC and NC stand for cotransporters with stoichiometry 1K^+^:1Cl‾ or 1Na^+^:1Cl‾. The latter can be represented by a single protein, the thiazide-sensitive Na-Cl cotransporter (SLC12 family), or by coordinated operation of the exchangers Na/H, SLC9, and Cl/HCO3, SLC26 (Garcia-Soto and Grinstein, 1990). The using the model with single parameters for characterization of each ion pathways is quite sufficient for successful description of the homeostasis in real cells at a real accuracy of the current experimental data. Some readers of our previous publications have expressed doubt that using our tool it is possible to obtain a unique set of parameters that provide an agreement between experimental and calculated data. Our mathematical comments on this matter can be found in Yurinskaya et al., 2019 (Notes Added in Response to Some Readers …).

The basic equations are presented below. Symbols and definitions used are shown in **Table 1**. Two mandatory conditions of macroscopic electroneutrality and osmotic balance:

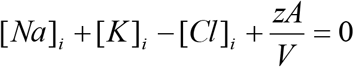

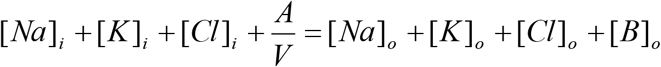

**Table 1.**
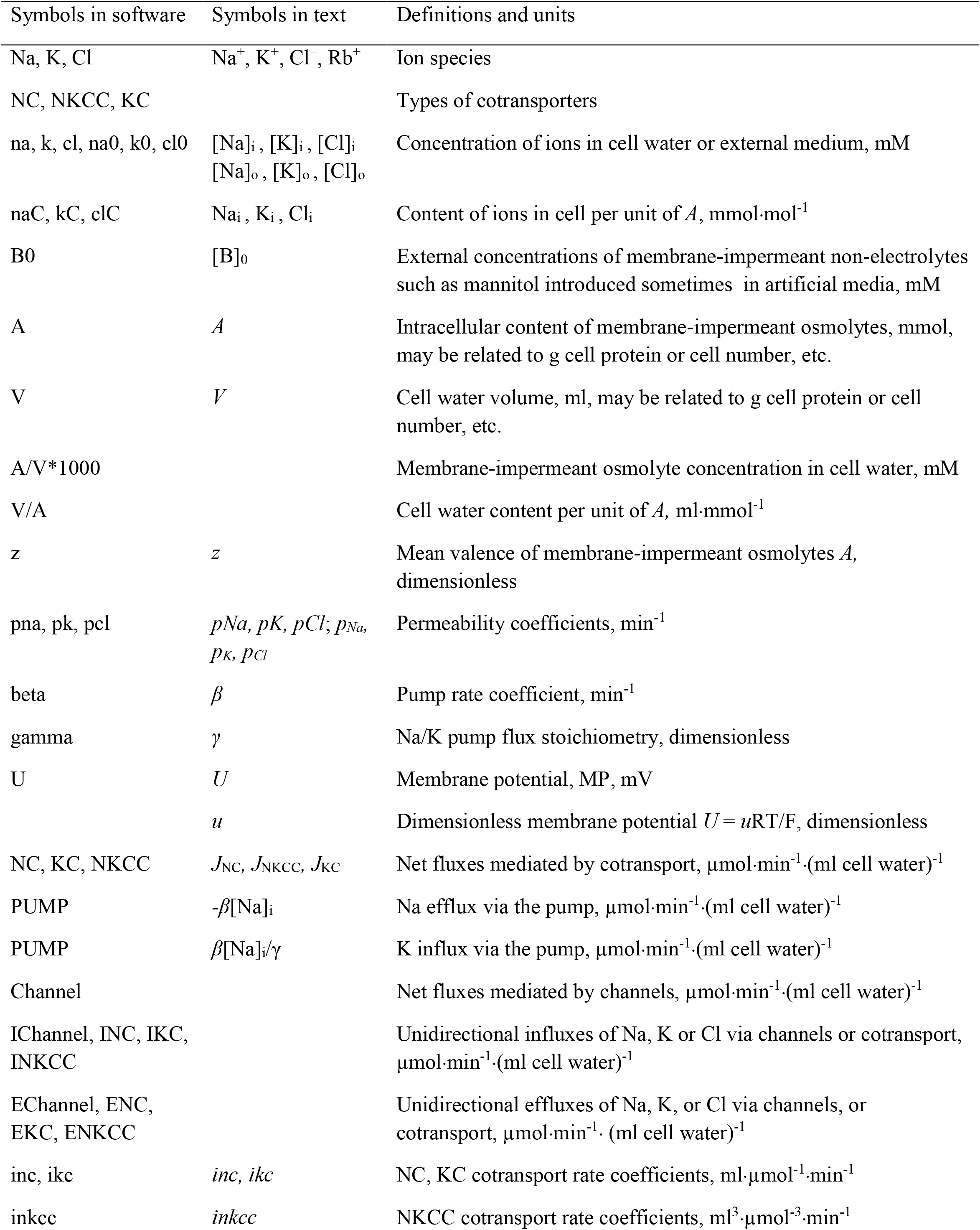
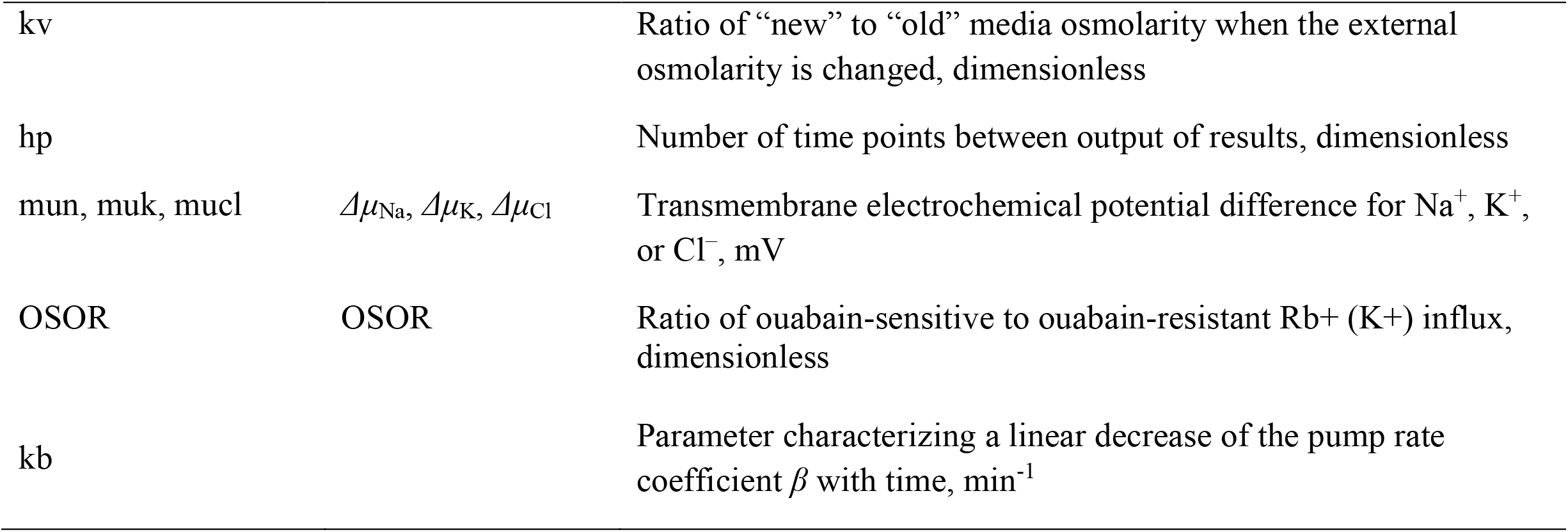
Symbols and definitions.

The flux equations:

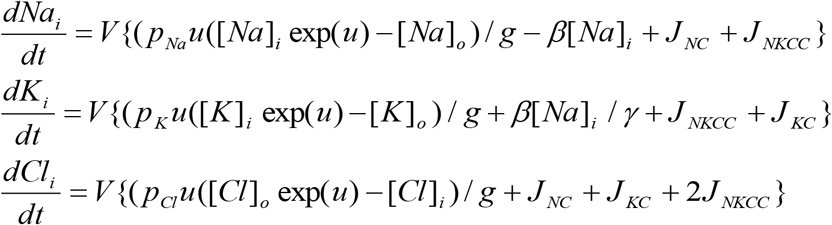

The left-hand sides of these three equations represent the rates of change of cell ion content. The right-hand sides express fluxes, where *u* is the dimensionless membrane potential related to the absolute values of membrane potential *U* (mV), as *U* = *u*RT/F = 26.7*u* for 37 °C and *g* = 1 − exp(*u*). The rate coefficients *p*_*Na*_, *p*_*K*_, *p*_*Cl*_ characterizing channel ion transfer are similar to the Goldman’s coefficients. Fluxes *J*_*NC*_, *J*_*KC*_, *J*_*NKCC*_ depend on internal and external ion concentrations as

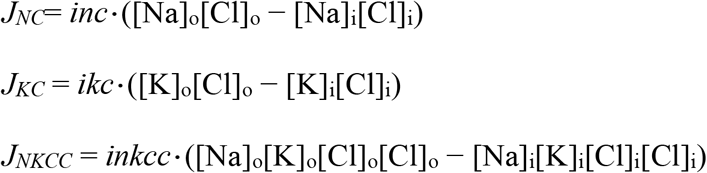

Here *inc, ikc*, and *inkcc* are the rate coefficients for cotransporters.

Transmembrane electrochemical potential differences for Na^+^, K^+^, and Cl^−^ were calculated as: *Δμ*_Na_ =26.7·ln([Na]i /[Na]o)+*U, Δμ*_K_ =26.7·ln([K]i /[K]o)+*U*, and*Δμ*_Cl_ =26.7·ln([Cl]i /[Cl]o)-*U*, respectively. The algorithm of the numerical solution of the system of these equations is considered in detail in (Vereninov et al., 2014), the using of the executable file is illustrated more in (Yurinskaya et al., 2019). The problems in determination of the multiple parameters in a system with multiple variables like cell ionic homeostasis are discussed in more detail in (Yurinskaya et al., 2019, 2020). The program BEZ02BC used in current study differs slightly from the previous program BEZ01B by replacing in the output table the columns prn, prk, prcl (time derivatives of concentrations) with the columns naC, kC, and clC representing intracellular content of Na^+^, K^+^, and Cl‾. Executable file to the program code BEZ01BC and Instruction: How to use programme code BEZ01BC.zip. are attached to the article electronic version.

## RESULTS

### 1. Observed and predicted changes in ionic homeostasis of U937 cells after stopping the Na/K pump, calculated for a system with cotransporters NC, KC, and NKCC and parameters like in U937 cells under a normal balanced state

The Na/K pump of the cell membrane is a key element of the cellular apparatus for holding the water balance of the animal cell, the dynamic balance in continuous movement K^+^, Na^+^ and Cl‾ between the exterior and cytoplasm, electrochemical gradients of these ions on the cell membrane and cell membrane potential (Cellular physiology and neurophysiology. 2nd ed., Elsevier Inc. 2012. 337 p). The dynamics of changes in the ionic homeostasis of the cell after stopping the pump has so far been studied using models with a limited list of cotransporters and without proper connection with experimental data. Our previous studies showed that computation based on the simplest model of cell ionic homeostasis including only the pump, Na^+^, K^+^, Cl‾ channels and cotransport NC predicts well the real-time dynamics of changes in ion concentrations and cell water content after blocking the pump even if the constant parameters of channels and NC cotransporter found for normal resting cells were used in calculations (Vereninov et al., 2014, 2016). However, we did not then consider other cation-chloride cotransporters. Now the calculations were carried out considering a various set of cotransporters.

Since the calculated parameters of channels and cotransporters depend on the measured characteristics differing due to the inevitable variability of the cells, **Table 2** shows two examples of U937 normal cells with different measured characteristics. The pump blocking experiment with ouabain and all related calculations were performed for cells *A*. There have been revealed certain differences in behavior of the model only with the NC cotransporter, which is required for most cells, and with a set of NC+NKCC, NC+KC, and NC+NKCC+KC cotransporters (**Figure 1**). The most significant differences appear with the addition of the NKCC cotransporter. A decrease in the water content in cells with the NKCC occurs with an extremum of 2 h. The possible delay in cell swelling, in this case, is caused by an increase in driving force moving Na^+^, K^+^, and Cl‾ via NKCC cotransporter at the stage when Na_i_ is increased significantly while K_i_ drops. Unfortunately, the discussed decrease is too small to be verified using current methods of analysis of cell water. In general, new calculations carried out with a wider list of cotransporters confirm that our computational approach allows us to quantitatively predict in real-time the dynamics of changes in the cellular ion and water homeostasis caused by stopping the pump. The accuracy of this prediction is at the level of the accuracy of the currently available experimental data. Good agreement between the predicted and the real behaviors of cells under stopping the pump shows that the approach used in the calculations is trustworthy. It should be emphasized that there were no parameters fitting in this case, as is often in modeling-simulation. All parameters used were determined outside the studied area and correspond to the normal unaltered cells.

**Table 2.** Basic characteristics of ion distribution measured for two examples of normal resting U937 cells (*A* and *B*), equilibrated with standard RPMI medium, and calculated for cells with different set of cotransporters.

|  | Measured characteristics, concentrations in mM |  |  |  |  |  |  |  |
| --- | --- | --- | --- | --- | --- | --- | --- | --- |
|  | A. Cells used in pump blocking experiments |  |  |  | B. Cells used in study of RVD |  |  |  |
|  | [K]<sub>i</sub> | [Na]<sub>i</sub> | [Cl]<sub>i</sub> | A<sup>z</sup> | [K]<sub>i</sub> | [Na]<sub>i</sub> | [Cl]<sub>i</sub> | A<sup>z</sup> |
|  | 156 | 35 | 70 | 49 | 147 | 38 | 45 | 80 |
|  | *Beta* 0.039, *z*-2.47, OSOR 3.06 |  |  |  | *Beta* 0.039, *z*-1.75, OSOR 3.89 |  |  |  |
|  | Supposed cotransporters and their parameters required for the balanced state |  |  |  |  |  |  |  |
| Parameters | NC+<br>KC+NKCC | NC+<br>NKCC | NC+KC | NC | NC+<br>KC+NKCC | NC+<br>NKCC | NC+KC | NC |
| *inc* | 7E-5 | 4.7E-5 | 5E-5 | 3E-5 | 7E-5 | 3E-5 | 4.87E-5 | 3E-5 |
| *ikc* | 3E-5 | — | 3E-5 | — | 8E-5 | — | 6E-5 | — |
| *inkcc* | 8E-9 | 8E-9 | — | — | 8E-9 | 7E-9 | — | — |
|  | Channel parameters computed for the selected cotransporters |  |  |  |  |  |  |  |
| *pna* | 0.0019 | 0.0032 | 0.0021 | 0.00317 | 0.0017 | 0.0043 | 0.00263 | 0.00382 |
| *pk* | 0.01 | 0.0149 | 0.0185 | 0.023 | 0.0115 | 0.0175 | 0.0165 | 0.022 |
| *pcl* | 0.004 | 0.00433 | 0.0029 | 0.00354 | 0.011 | 0.0139 | 0.006 | 0.0091 |
|  | Computed homeostasis characteristics for the selected cotransporters, mV |  |  |  |  |  |  |  |
| *U* | -45.2 | -42.0 | -55.5 | -50.5 | -45.0 | -37.6 | -49.3 | -44.7 |
| *mucl* | +31.7 | +28.6 | +42.0 | +37.0 | +19.8 | +12.3 | +24.0 | +19.4 |
| *mun* | -82.2 | -79.1 | -92.5 | -87.5 | -79.9 | -72.4 | -84.1 | -79.5 |
| *muk* | +42.7 | +45.8 | +32.4 | +37.4 | +41.3 | +48.7 | +37.0 | +41.6 |

**Figure 1.**
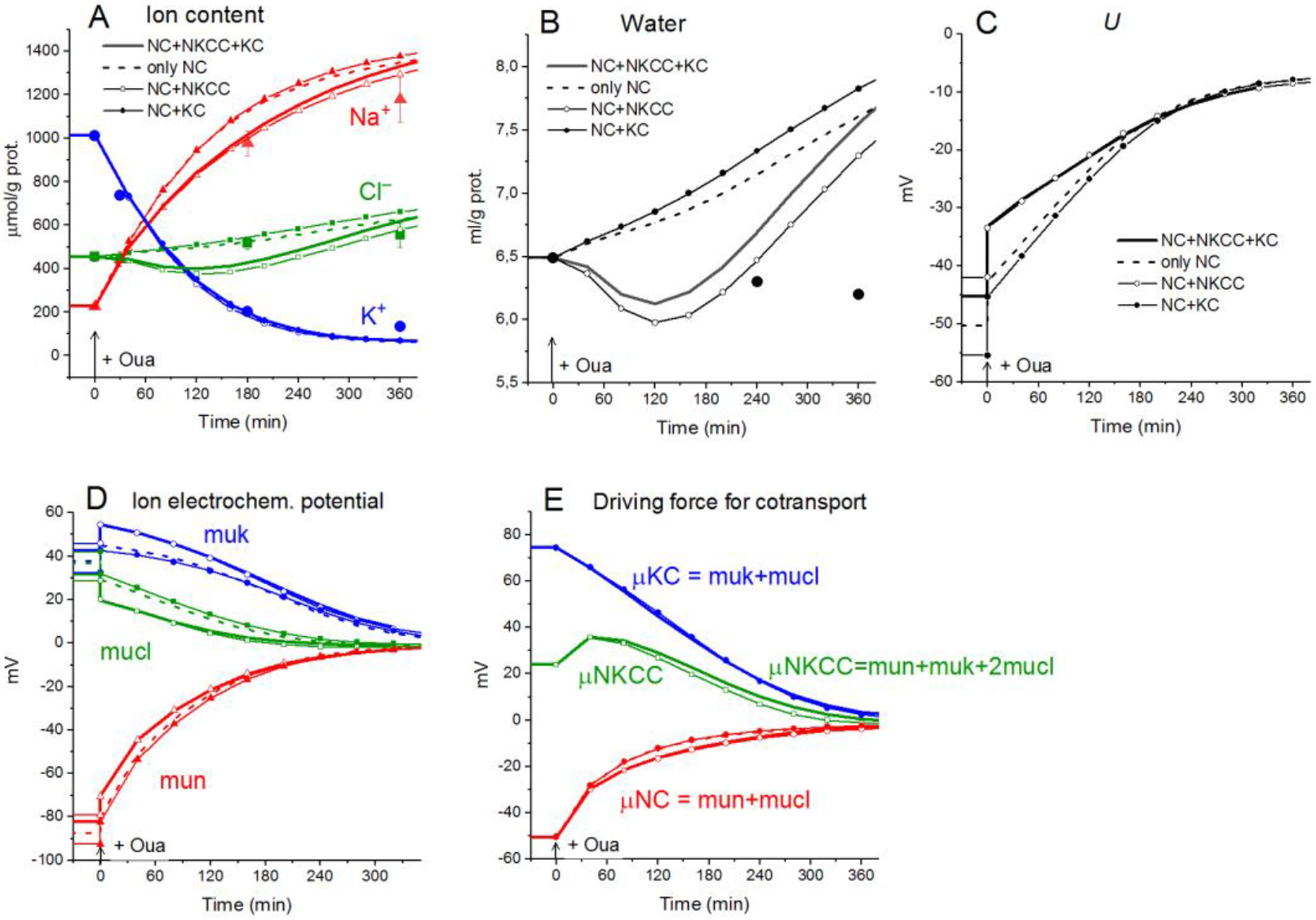
**Disturbance of cell ionic homeostasis after stopping the pump in the real U937 cells (large symbols) and in U937 cell model with different set of cotransporters at the unchanged over time parameters as in U937 cells, equilibrated with the standard RPMI medium (lines with or without small symbols).** The pump was blocked by ouabain at t = 0, corresponding to a change in *beta* from 0.039 to 0. The other parameters were unchanged and are shown in Table 2, cells *A*. Water in ml/g protein was obtained by multiplying the calculated V/A by the content of impermeant osmolytes *A* in mol/g protein.

Cell swelling after blocking the pump deserves additional remarks. It is known that a cell with a membrane permeable to water and external ions and impermeable to some intracellular ions behaves like a Donnan system (Hoffman et al., 2009; Jentsch, 2016; Delpire and Gagnon, 2018). The water balance between solutions separated by membrane cannot physically be achieved in such a system if the external medium contains only ions freely penetrating through the membrane. The concentration of anions inside the real cells, impermeant through the plasma membrane and represented mostly by proteins, nucleic acids and organic and inorganic phosphates is equal to the difference between the concentrations of intracellular Cl‾ and the sum of intracellular K^+^ and Na^+^, i.e., significant. In an environment with ions and uncharged osmolytes that freely penetrate the membrane, the cells, after stopping the pump, must infinitely swell until the membrane ruptures. The first function of the Na/K pump is the preventing the Donnan’s water disbalance. Pumping sodium out of the cell makes the membrane virtually impermeable to sodium and this ion behaves as an impermeant external cation. The Donnan effect due to the quasi-impermeant extracellular sodium balances the Donnan effect caused by the impermeant intracellular anions. Hence, cell behaves as the double-Donnan system (see using the term in Freedman and Hoffman, 1979; Fraser and Huang, 2007). A quantitative study of changes in ionic and water homeostasis in U937 cells after blocking the pump with ouabain and mathematical modeling of these changes in real experiments shows that the system eventually comes to a new balanced state if the medium contains at least a small concentration of impermeant osmolytes. In our calculation in case of the RPMI medium B0 is equal to 48 mM (**Table 3, Figure 2)**. The final water level in U937 cells in a standard medium RPMI with ouabain is 1.4 times higher than the initial and fully meets the balance criteria (**Figure 2)**. Hence, the 48 mM concentration of impermeant charged osmolytes B in the real physiological media is sufficient to prevent unlimited cell swelling in our case, but the system degradation is significant. The question “Why real cells do not swell infinitely after blocking the pump” has many answers. It is not only due to a parallel alteration of electroconductive channels, as we wrote earlier (Yurinskaya et al., 2011, 2020). The swelling is highly retarded because of decreasing of the driving forces for all ion pathways (**Figure 1 D, E**). Finally, some external impermeant osmolyte is always present in experiments with real cells. This is not an idle question, because it is the Donnan effect that leads to an extremely dangerous cerebral edema in brain ischemia when the sodium pump stops due to the ATP deficiency (Dijkstra et al., 2016; Okada al., 2019). Osmolysis of human RBC infected by the malaria plasmodium plays significant role in a pathology of this disease (Mauritz et al., 2009; Waldecker et al., 2017).

**Table 3.** Dynamics of the net and unidirectional K^+^, Na^+^, and Cl^−^ fluxes after stopping the pump calculated for the model with all main cotransporters at the unchanged over time parameters as in U937 cells, equilibrated with the standard RPMI medium: Na_o_ 140, K_o_ 5.8; Cl_o_ 116 mM, B_o_ 48.2 mM

| Ion | Time, min | μ, mV | Net fluxes, total | Unidirectional fluxes |  |  |  |  |  |  |  |
| --- | --- | --- | --- | --- | --- | --- | --- | --- | --- | --- | --- |
|  |  |  |  | Influxes |  |  |  | Effluxes |  |  |  |
|  |  |  |  | IChannel | PUMP | IKC | INKCC | EChannel | PUMP | EKC | ENKCC |
| K<sup>+</sup> | 0 | 42.7 | 0.0000 | 0.1203 | 0.9098 | 0.0202 | 0.0874 | -0.5956 | 0 | -0.3278 | -0.2142 |
|  | 30 | 52.0 | -1.0538 | 0.0959 | 0 | 0.0202 | 0.0874 | -0.6733 | 0 | -0.2580 | -0.3259 |
|  | 48 | 50.0 | -0.9646 | 0.0934 | 0 | 0.0202 | 0.0874 | -0.6069 | 0 | -0.2193 | -0.3393 |
|  | 240 | 17.5 | -0.1247 | 0.0722 | 0 | 0.0202 | 0.0874 | -0.1387 | 0 | -0.0382 | -0.1275 |
|  | 480 | 1.3 | -0.0016 | 0.0660 | 0 | 0.0202 | 0.0874 | -0.0693 | 0 | -0.0208 | -0.0851 |
|  | 1200 | 0.0 | 0.0000 | 0.0646 | 0 | 0.0202 | 0.0874 | -0.0646 | 0 | -0.0202 | -0.0873 |
|  | 3600 | 0.0 | 0.0000 | 0.0645 | 0 | 0.0202 | 0.0874 | -0.0645 | 0 | -0.0202 | -0.0874 |
|  |  |  |  | IChannel | INC | IKC | INKCC | EChannel | ENC | PUMP | ENKCC |
| Na<sup>+</sup> | 0 | -82.2 | 0.0000 | 0.5517 | 1.1368 | 0 | 0.0874 | -0.0254 | -0.1716 | -1.3647 | -0.2142 |
|  | 30 | -48.9 | 0.9363 | 0.4400 | 1.1368 | 0 | 0.0874 | -0.0704 | -0.3315 | 0 | -0.3259 |
|  | 48 | -41.2 | 0.8156 | 0.4283 | 1.1368 | 0 | 0.0874 | -0.0915 | -0.4061 | 0 | -0.3393 |
|  | 240 | -6.6 | 0.2934 | 0.3309 | 1.1368 | 0 | 0.0874 | -0.2587 | -0.8755 | 0 | -0.1275 |
|  | 480 | -1.0 | 0.0749 | 0.3026 | 1.1368 | 0 | 0.0874 | -0.2917 | -1.0751 | 0 | -0.0851 |
|  | 1200 | -0.0 | 0.0020 | 0.2960 | 1.1368 | 0 | 0.0874 | -0.2958 | -1.1352 | 0 | -0.0873 |
|  | 3600 | -0.0 | 0.0000 | 0.2959 | 1.1368 | 0 | 0.0874 | -0.2959 | -1.1368 | 0 | -0.0874 |
|  |  |  |  | IChannel | INC | IKC | INKCC | EChannel | ENC | EKC | ENKCC |
| Cl<sup>-</sup> | 0 | 31.7 | 0.0000 | 0.1772 | 1.1368 | 0.0202 | 0.1748 | -0.5811 | -0.1716 | -0.3278 | -0.4285 |
|  | 30 | 16.0 | -0.1178 | 0.2535 | 1.1368 | 0.0202 | 0.1748 | -0.4618 | -0.3315 | 0.2580 | -0.6517 |
|  | 48 | 13.7 | -0.1494 | 0.2635 | 1.1368 | 0.0202 | 0.1748 | -0.4406 | -0.4061 | -0.2193 | -0.6786 |
|  | 240 | -0.4 | 0.1686 | 0.3666 | 1.1368 | 0.0202 | 0.1748 | -0.3611 | -0.8755 | -0.0382 | -0.2550 |
|  | 480 | -0.5 | 0.0733 | 0.4055 | 1.1368 | 0.0202 | 0.1748 | -0.3978 | -1.0751 | -0.0208 | -0.1703 |
|  | 1200 | -0.0 | 0.0020 | 0.4153 | 1.1368 | 0.0202 | 0.1748 | -0.4151 | -1.1352 | -0.0202 | -0.1747 |
|  | 3600 | -0.0 | 0.0000 | 0.4155 | 1.1368 | 0.0202 | 0.1748 | -0.4155 | -1.1368 | -0.0202 | -0.1748 |

**Figure 2.**
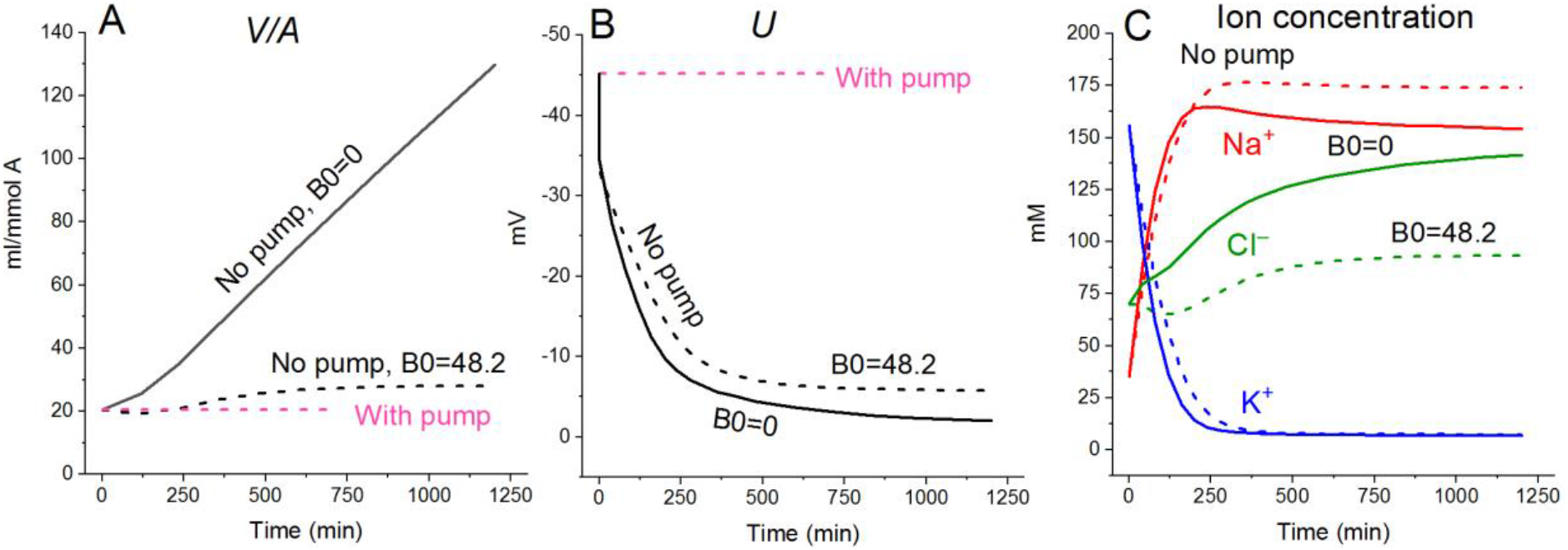
**Effect of impermeant external osmolytes B_o_ on cell swelling caused by blocking the pump calculated for the U937 cell model with a full set of cotransporters (NC+ KC+ NKCC) at the unchanged over time parameters as in U937 cells, equilibrated with standard RPMI medium.** The calculation was carried out for B_o_=0 (solid lines, Na_o_ 149.2, K_o_ 5.8, Cl_o_ 155 mM) or for standard medium (dashed lines, Na_o_ 140, K_o_ 5.8; Cl_o_ 116 mM, B_o_ 48.2 mM). The pump was blocked at t = 0 by changing the *beta* from 0.039 to 0 (“No pump” in the figure). The other parameters remain constant and are shown in Table 2, cells *A* with a full set of cotransporters.

### 2. Cotransporters in rearrangement of ionic homeostasis in cells transferred into hypoosmolar media

#### 2.1. Regulatory volume decrease (RVD) in the model with all main cotransporters at the unchanged over time parameters like in U937 cells equilibrated with standard RPMI

Water penetrates through the cell membrane more easily than ions, and after replacing the normal medium with a hypoosmolar solution, the water content in the cell increases sharply according to the well-known law of the water-osmotic balance of the cell. Changes in intracellular ion concentration, membrane potential, and electrochemical ion gradients across the cell membrane also occur rapidly, while changes in ion content take time. The phenomenon with the abbreviation RVD (Regulatory Volume Decrease), observed in most cells as a response to a decrease in osmolarity of the medium, has long attracted the attention of many researchers.

However, the calculation of the behavior of such a complex multi-parameter system as the electrochemical system of a cell still presents significant difficulties. We believe that our software can be useful for studying the physical basis of ion flux balance rearrangement during RVD. Analysis of RVD is demonstrated here on U937 cells because we have a set of necessary parameters for these cells that were obtained in our own experiments (**Table 2)**.

First, the calculation shows that a balanced state in the distribution of monovalent ions is established in cells placed in a hypoosmolar medium over time **(Figure 3)**. Calculation of fluxes confirms that this is a truly balanced state and that the total net fluxes of each species of ions decline with time to zero, even though the ionic electrochemical gradients remain nonzero (**Table 4**). Due to zero net fluxes, the ion content in the cell ceases to change and a water balance is established. The influx and efflux via each individual pathway remain in this case unbalanced but the integral net flux via all pathways becomes zero. The next important point is that in the cases under consideration, there is a time-dependent decrease in the volume of cells by the type of the physiological RVD without any specific “regulatory” changes in membrane channels and transporters (**Figure 3**). Therefore, changes in ionic homeostasis and fluxes observed in real cells during the transition to a hypoosmolar medium, even those affected by specific inhibitors, cannot serve as evidence that specific “regulatory” changes are triggered in the corresponding pathway in a hypoosmolar medium, as it is usually assumed.

**Table 4.** **Dynamics of the net and unidirectional fluxes of K^+^, Na^+^, and Cl^-^during transition from normal medium 310 mOsm to the hypoosmolar solution 160 mOsm, calculated for the system with the complete set of cotransporters at the unchanged over time parameters as in U937 cells equilibrated with standard RPMI medium.** Parameters are shown in Table 2 for Cells *B*, NC+KC+NKCC. Fluxes are given in µmol min^−1^(ml cell water)^−1^. An initial increase and following decrease in the K^+^ net flux with a local extremum at the 24 min are marked in red.

| Ion | Time<br>in160<br>mOsM,<br>min | Ion<br>electrochemical<br>gradients, mV | Net<br>fluxes,<br>total | Unidirectional fluxes |  |  |  |  |  |  |  |
| --- | --- | --- | --- | --- | --- | --- | --- | --- | --- | --- | --- |
|  |  |  |  | Influxes |  |  |  | Effluxes |  |  |  |
| K+ | Before | *muk*<br>41.3 | 0.0000 | PUMP<br>0.9880 | IChannel<br>0.1381 | IKC<br>0.0538 | INKCC<br>0.0874 | EChannel<br>-0.6477 | PUMP<br>0 | EKC<br>-0.5291 | ENKCC<br>-0.0905 |
|  | 1 | 28.5 | 0.1317 | 0.5010 | 0.1292 | 0.0190 | 0.0051 | -0.3759 | 0 | -0.1405 | -0.0062 |
|  | 24 | 29.1 | -0.0013 | 0.3664 | 0.1319 | 0.0190 | 0.0051 | -0.3924 | 0 | -0.1278 | -0.0035 |
|  | 48 | 28.4 | -0.0395 | 0.3088 | 0.1350 | 0.0190 | 0.0051 | -0.3906 | 0 | -0.1145 | -0.0023 |
|  | 60 | 27.9 | -0.0427 | 0.2950 | 0.1365 | 0.0190 | 0.0051 | -0.3873 | 0 | -0.1090 | -0.0020 |
|  | 120 | 25.8 | -0.0233 | 0.2759 | 0.1412 | 0.0190 | 0.0051 | -0.3710 | 0 | -0.0922 | -0.0013 |
|  | 240 | 24.6 | -0.0027 | 0.2757 | 0.1437 | 0.0190 | 0.0051 | -0.3609 | 0 | -0.0843 | -0.0011 |
|  | 480 | 24.4 | 0.0000 | 0.2760 | 0.1440 | 0.0190 | 0.0051 | -0.3596 | 0 | -0.0834 | -0.0011 |
| Na+ | Before | *mun*<br>-79.9 | 0.0000 | IChannel<br>0.4927 | INC<br>1.1368 | PUMP<br>0 | INKCC<br>0.0874 | PUMP<br>-1.4820 | ENC<br>-0.1197 | EChannel<br>-0.0248 | ENKCC<br>-0.0905 |
|  | 1 | -72.7 | -0.3971 | 0.2140 | 0.1866 | 0 | 0.0051 | -0.7515 | -0.0311 | -0.0140 | -0.0062 |
|  | 24 | -82.6 | -0.1520 | 0.2186 | 0.1866 | 0 | 0.0051 | -0.5496 | -0.0191 | -0.0099 | -0.0035 |
|  | 48 | -88.8 | -0.0721 | 0.2237 | 0.1866 | 0 | 0.0051 | -0.4632 | -0.0139 | -0.0080 | -0.0023 |
|  | 60 | -90.8 | -0.0468 | 0.2261 | 0.1866 | 0 | 0.0051 | -0.4425 | -0.0126 | -0.0075 | -0.0020 |
|  | 120 | -95.1 | -0.0059 | 0.2339 | 0.1866 | 0 | 0.0051 | -0.4138 | -0.0098 | -0.0066 | -0.0013 |
|  | 240 | -96.4 | -0.0003 | 0.2381 | 0.1866 | 0 | 0.0051 | -0.4135 | -0.0089 | -0.0064 | -0.0011 |
|  | 480 | -96.5 | 0.0000 | 0.2386 | 0.1866 | 0 | 0.0051 | -0.4139 | -0.0088 | -0.0064 | -0.0011 |
| Cl- | Before | *mucl*<br>19.8 | 0.0000 | IChannel<br>0.4889 | INC<br>1.1368 | IKC<br>0.0538 | INKCC<br>0.1748 | EChannel<br>-1.0246 | ENC<br>-0.1197 | EKC<br>-0.5291 | ENKCC<br>-0.1809 |
|  | 1 | 24.9 | -0.2656 | 0.1934 | 0.1866 | 0.0190 | 0.0101 | -0.4906 | -0.0311 | -0.1405 | -0.0125 |
|  | 24 | 21.7 | -0.1732 | 0.1867 | 0.1866 | 0.0190 | 0.0101 | -0.4216 | -0.0191 | -0.1278 | -0.0070 |
|  | 48 | 19.6 | -0.1114 | 0.1795 | 0.1866 | 0.0190 | 0.0101 | -0.3736 | -0.0139 | -0.1145 | -0.0046 |
|  | 60 | 18.8 | -0.0895 | 0.1763 | 0.1866 | 0.0190 | 0.0101 | -0.3560 | -0.0126 | -0.1090 | -0.0039 |
|  | 120 | 16.4 | -0.0292 | 0.1661 | 0.1866 | 0.0190 | 0.0101 | -0.3065 | -0.0098 | -0.0922 | -0.0026 |
|  | 240 | 15.2 | -0.0030 | 0.1610 | 0.1866 | 0.0190 | 0.0101 | -0.2843 | -0.0089 | -0.0843 | -0.0021 |
|  | 480 | 15.0 | 0.0001 | 0.1604 | 0.1866 | 0.0190 | 0.0101 | -0.2818 | -0.0088 | -0.0834 | -0.0021 |

**Figure 3.**
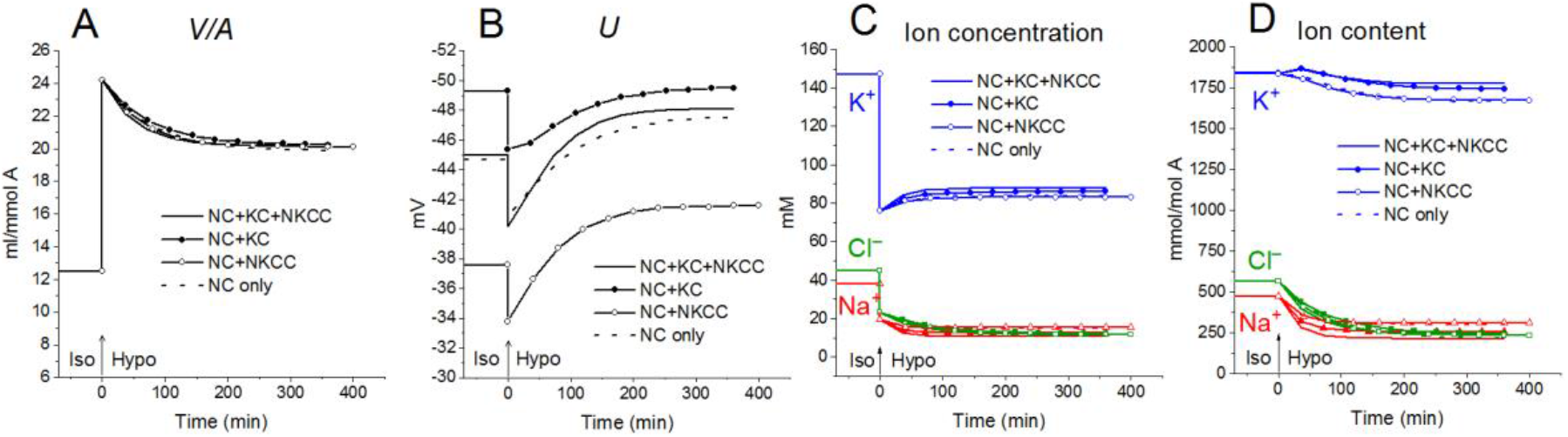
**Rearrangement of ionic homeostasis during the Iso-Hypo transition, calculated for a system with different sets of cotransporters at the unchanged over time parameters like in U937 cells, equilibrated with the standard RPMI medium.** The parameters for the calculation are given in Table 2, cells *B*. The arrows at t = 0 show the replacement of a standard medium of 310 mOsm with a hypoosmolar medium of 160 mOsm,obtained by decreasing the NaCl concentration by 75 mM.

Variations in the types of cotransporters in the cell membrane have no significant effect on the time course of changes in cell water during the Iso-Hypo transition **(Figure 3)**. Consequently, the type of cotransporters cannot be identified by studying changes in V/A. The dynamics of *U* is more dependent on carriers, even though they are “electroneutral”. This is a good example of the fact that the electroneutrality of the coupled transport of ions in a cotransporter does not mean that the cotransporter is “electroneutral” when operating in a complex system. Unfortunately, it is difficult to measure the difference of about 10 mV in the membrane potential of cells in a population with sufficient accuracy to identify cotransporters by this way. The difference in the early increase in net K^+^ flux and intracellular K^+^ content depending on cotransporters is also small and difficult to accurately measure.

A hypoosmolar medium is usually prepared by diluting the normal medium or excluding some of the NaCl from it. A decrease in external NaCl concentration is the essential factor which changes the forces driving ions through the plasma membrane and leads to disturbance of a balance of ion fluxes across the cell membrane. To separate the roles of a decrease in the external concentration of NaCl and a decrease in the osmolarityof the medium, three schemes were calculated (**Figure 4**): (1) a simple decrease in NaCl (curve 1), (2) a decrease in external NaCl by 75 mM compensated by addition of equimolar 150 mM sucrose, and (3) a variant when hypoosmolar solution is prepared by excluding of 150 mM sucrose from a medium of 310 mOsm, initially containing 75 mM NaCl and 150 mM sucrose (curve 3). No change in external NaCl concentration occurs in the last case. Thus, in our modeling, two cellular responses, to a decrease in NaCl concentration and to a change in osmolarity, are generated independently. A decrease in the NaCl concentration is followed by RVD in both iso- and in hypotonic solution **(Figure 4**, curves 1 and 2). A decrease in extracellular osmolarity due to the exclusion of 150 mM external sucrose without changing the extracellular NaCl concentration is not accompanied by RVD (**Figure 4**, curve 3).

**Figure 4.**
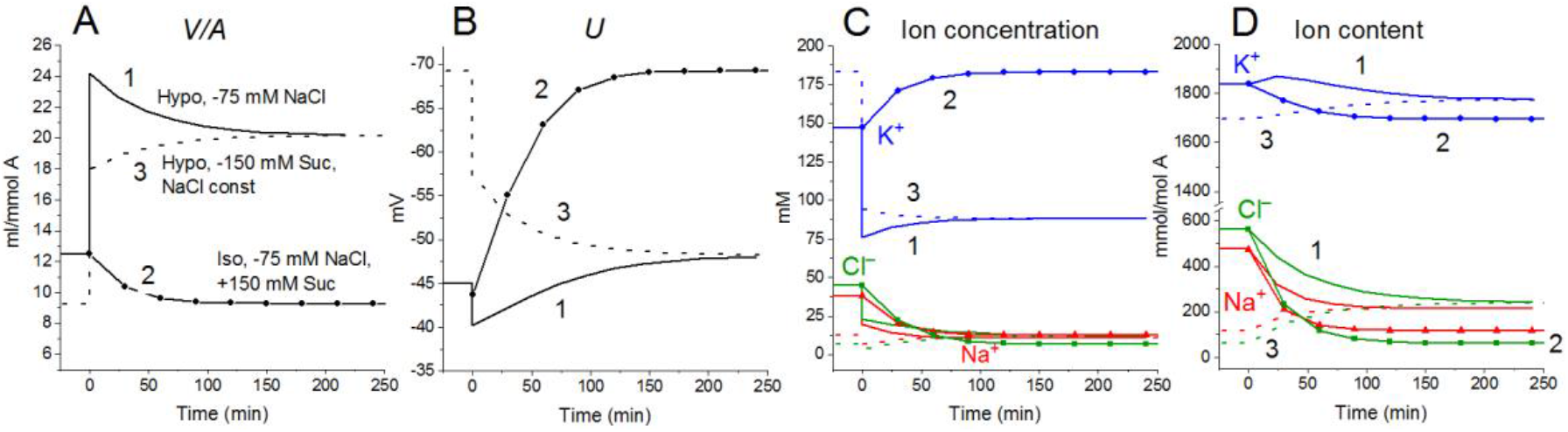
**Rearrangement of ionic homeostasis during transition to hypotonic medium with and without a decrease in NaCl concentration, calculated for a system with a full set of cotransporters (NC+ KC+ NKCC) at the unchanged over time parameters as in U937 cells, equilibrated with standard RPMI medium.** The standard medium was replaced at t=0 by (1) hypoosmolar medium of 160 mOsm obtained by removing 75 mM NaCl from standard isoosmolar medium, (2) isoosmolar medium of 310 mOsm, prepared by removing 75 mM NaCl and adding 150 mM sucrose instead, (3) a medium of 160 mOsm obtained by removing 150 mM sucrose from an isoosmolar medium without a decrease in NaCl concentration. The concentrations of Na^+^ and Cl^−^ in media (2) and (3) were 65 and 41 mM, respectively.

**Figure 5.**
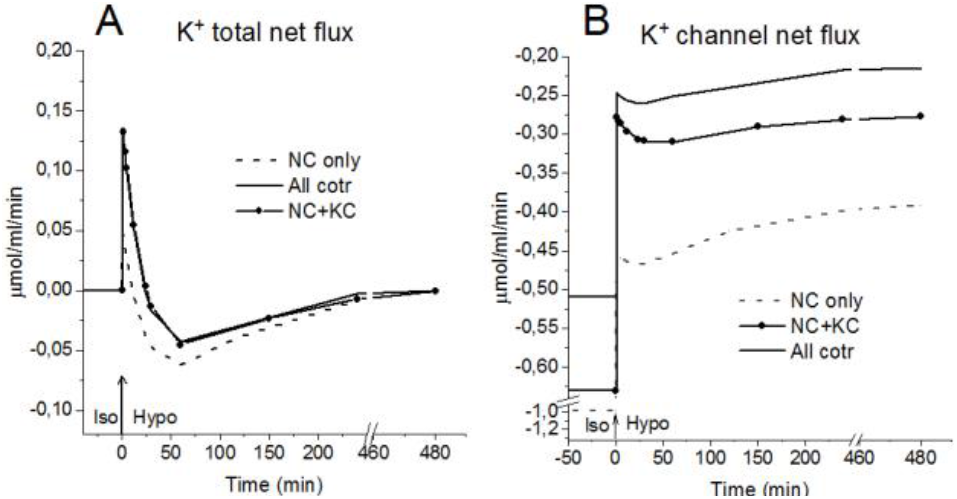
K^+^ fluxes during the Iso-Hypo transition, calculated for a system with different sets of cotransporters at the unchanged over time parameters as in U937 cells, equilibrated with standard RPMI medium. The arrows at t = 0 show the replacement of the standard 310 mOsm medium with a 160 mOsm hypoosmolar medium,obtained by decreasing the NaCl concentration by 75 mM.

An unexpected and interesting effect observed during the Iso-Hypo transition is an increase in the net flux of K^+^ into the cell at the initial stage of rearrangement of homeostasis, which then decreases, transforms into an outgoing net flux, which decreases to zero when the cell comes to new balanced state **(Figures 3-5, Table 4**). This effect, like the initial increase in intracellular K^+^ content, was intuitively impossible to expect. It can be explained by the different dependence of unidirectional fluxes via different pathways on changes in the intracellular concentration of ions in cells placed in a hypoosmolar medium, and the asynchrony of changes in partial fluxes. This example shows that non-monotonic changes in ion fluxes and, consequently, ion concentration during rearrangement of such a complex system as ionic homeostasis of the cell, can occur without specific changes in the channels and transporters of the cell membrane.

Rearrangement of ionic homeostasis caused by the hypoosmolar medium is reversible and is followed by an increase in volume during the reverse Hypo-Iso transition, which resembles the so-called RVI after RVD (Hoffmann et al., 2009). No specific alterations of channels and transporters are required for the model RVI at the transition Hypo-Iso. Changes in cell volume, in this case, are practically independent of the type of cotransporter, as in the direct Iso-Hypo transition, while the recovery curves of *U*, net K^+^ flux, and K^+^ content depend, although not strongly **(Figure 6)**. It will be shown below that decreasing NC rate coefficient simultaneously with the transition Iso-Hypo or Hypo-Iso changes RVD and RVI (**Figure 7**).

**Figure 6.**
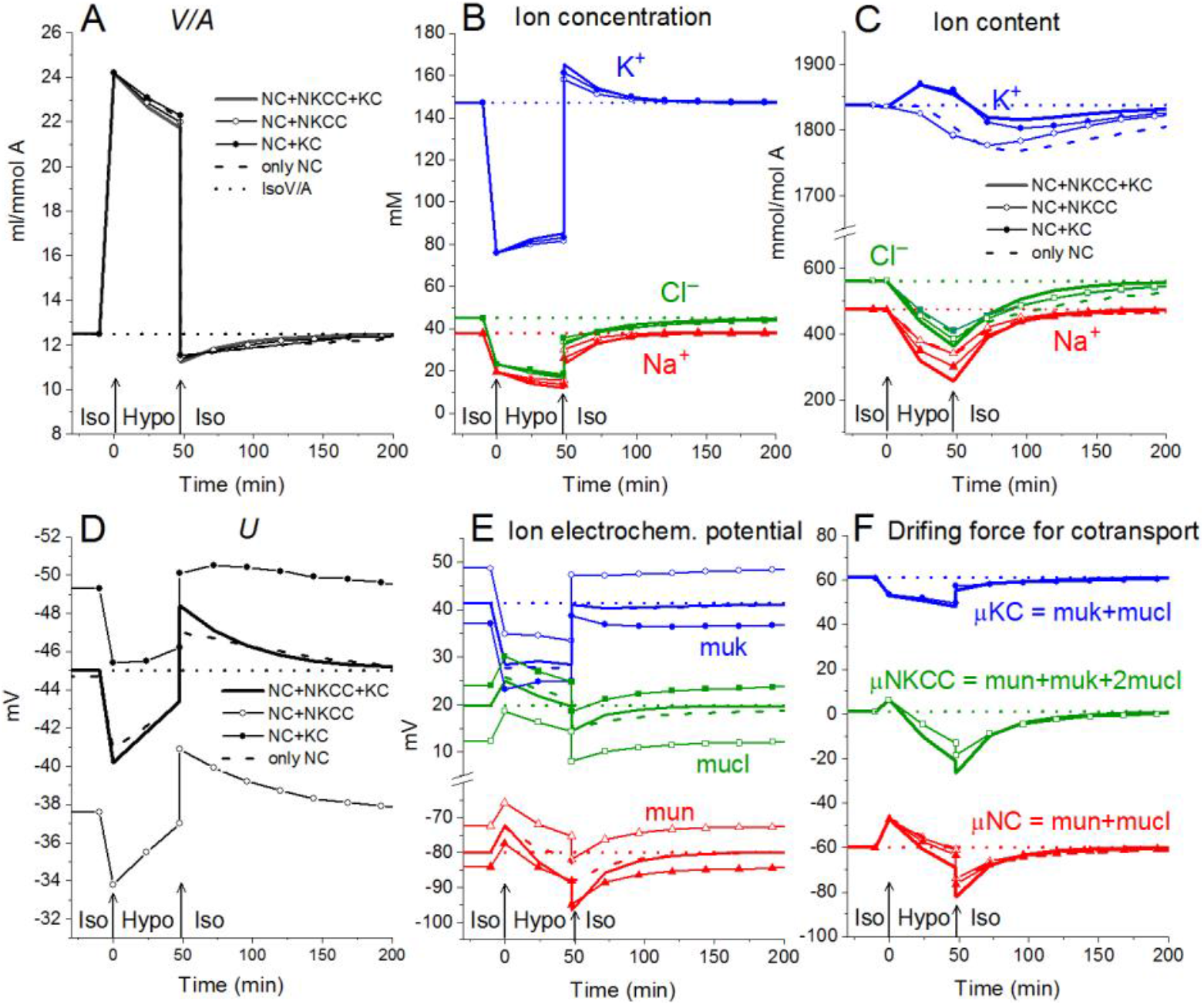
**Rearrangement of ionic homeostasis during the Iso-Hypo transition and the reverse transition to a normal medium, calculated for a system with different sets of cotransporters at the unchanged over time parameters as inU937 cells, equilibrated with standard RPMI medium.** A hypoosmolar medium of 160 mOsm was obtained by decreasing the NaCl concentration by 75 mM. The parameters for the calculation are given in Table 2, cells *B*.

**Figure 7.**
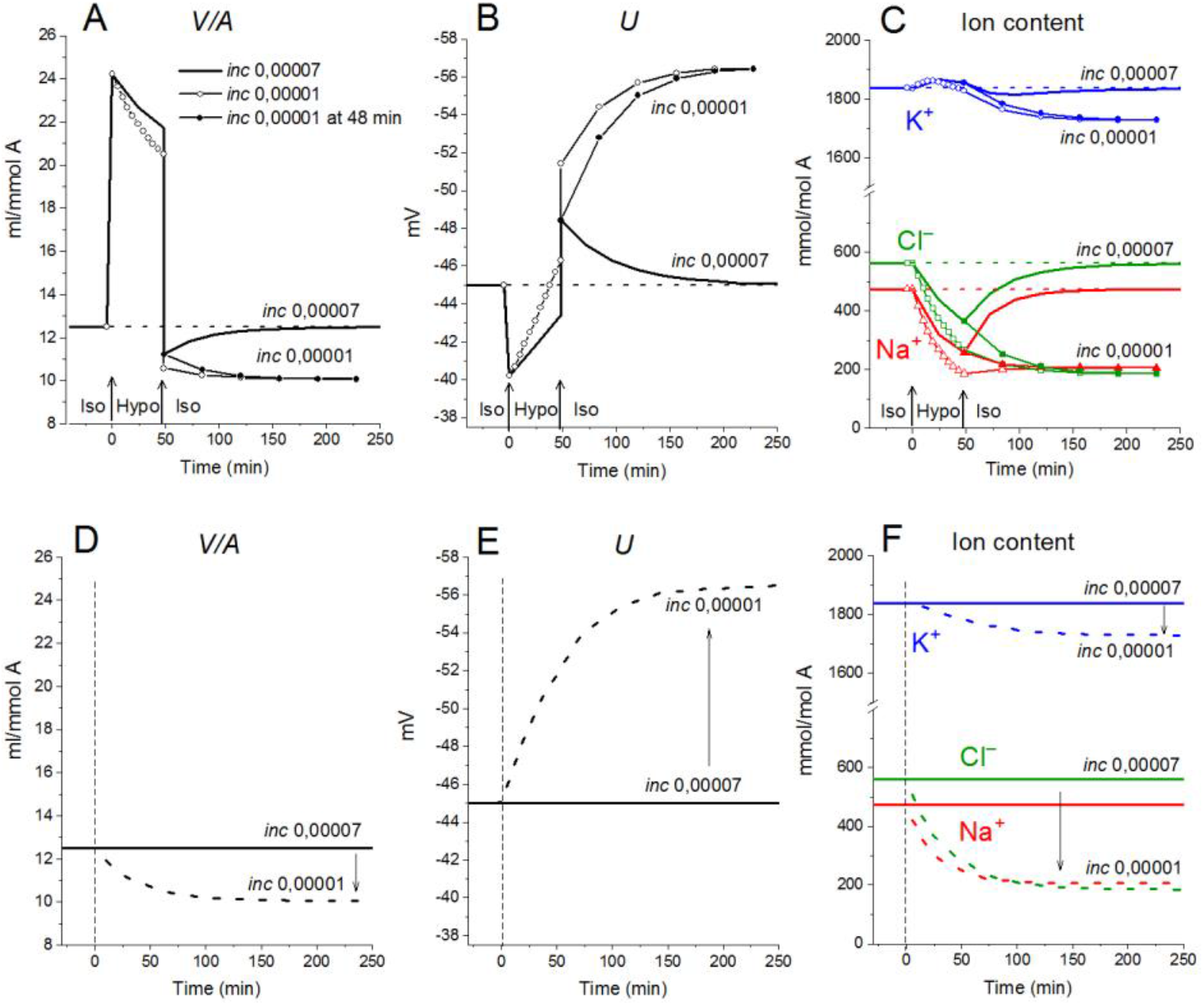
**The effect of NC rate constant on the rearrangement of ionic homeostasis during the Iso-Hypo-Iso transitions (A-C) and in a normal 310 mOsm medium without changes in external osmolarity (D-F), calculated for a system with a full set ofcotransporters (NC+ KC+ NKCC).** Changes in the *inc* coefficient are displayed on the graphs. (A-C) *inc* was changed either simultaneously with the transition to a hypotonic medium (at t=0) or upon a return to a normal isotonic (at t=48 min).Other parameters remain unchanged.

#### 2.2. Rearrangement of ionic homeostasis in the U937 cell model due to changing membrane parameters in normal and hypoosmolar media

Although a response resembling physiological RVD can be seen in the physical system without changes in the properties of channels and cotransporters, many experimental data shows that these changes occur in living cells placed in a hypoosmolar environment (Hoffmann et al 2009; Delpire, Gagnon, 2018). Modeling can help to quantify the relationship between RVD and alteration of channels and transporters in a hypoosmolar environment. When the properties of channels and carriers change simultaneously with the transition to a hypoosmolar medium, two effects are summed up: one is associated with the Iso-Hypo transition, and the other is associated with a change in channels and transporters. They can be distinguished only by simulation which shows that the effects of changes in membrane parameters without changes in external osmolarity are significant.

##### 2.2.1. Increasing the rate coefficients pK, pCl and ikc enhances RVD in hypoosmolar medium

The activation of the K^+^ and Cl‾ channels in cells placed into hypoosmolar media is believed to be important in reducing the intracellular levels of K^+^ and Cl‾, which underlie RVD. Indeed, an increase in pK and pCl results in a time-dependent decrease in cell volume (**Figure 8 A)**. Similar decrease can be caused by a decrease in the NC rate coefficient (**Figure 8 E)**. Changes in the ion content in these cases differ significantly, as well as the changes in the membrane potential *U*. Modeling shows that the considered effects of channel permeability and NC depend on the basic set of cotransporters in the membrane and that there may be unpredictable phenomena in the behavior of the system, for example, the K^+^ content may decrease monotonically or initially pass through a maximum or minimum, depending on the conditions (**Figure 8)**. A simultaneous change in pK and pCl can reduce the amount of water in cells more than their change separately, but the cell volume is restored even in this case by only half. Variation of the KC rate coefficients (*ikc*) affects the ionic homeostasis of the cell in almost the same way as the change in pK (data not shown).

**Figure 8.**
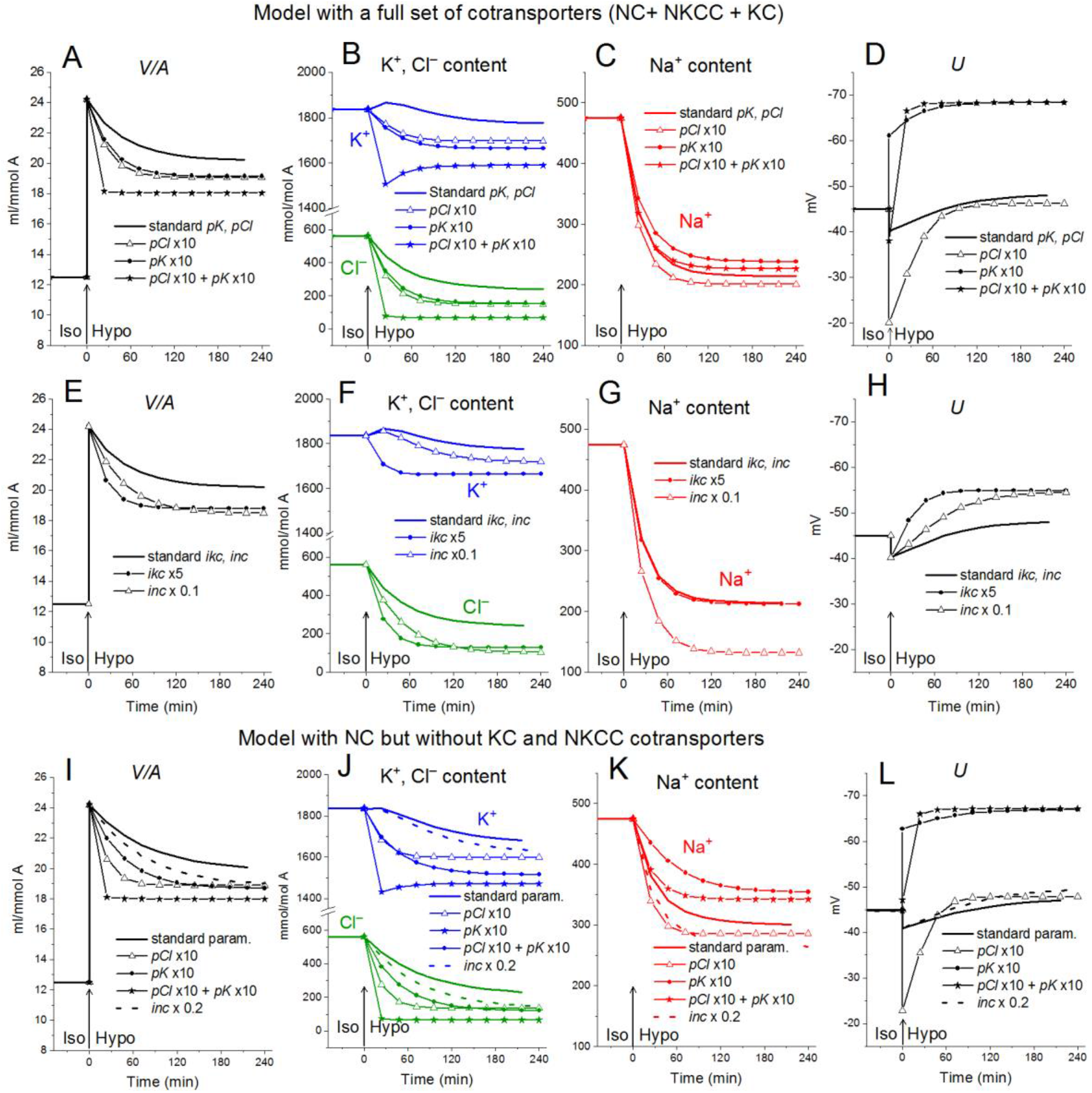
The effect of pK, pCl, NC and KC rate constant on the rearrangement of ionic homeostasis in the Iso-Hypo transition, calculated for the U937 cell model with a full set of cotransporters and for the model with only NC. The parameters of membrane transport changed simultaneously with the transition to a hypotonic medium. Changes in one of the parameters are indicated on the graphs. The rest of the parameters remain unchanged.

##### 2.2.2. Decreasing the NC rate coefficient enhances RVD

Decreasing the NC rate coefficient (*inc*) in hypoosmolar medium with a decreased NaCl enhances RVD (**Figures 7 A, 8 E-L**). It should be noted here that under isoosmolar conditions NC affects the ionic homeostasis of the cell in the same way as in the hypoosmolar medium (**Figure 7 D**). Interestingly, decreasing *inc* declines RVI observed at the transfer of cells from hypoosmolar to isoosmolar medium (so-called RVI-after-RVD).

##### 2.2.3. Changes in partial fluxes underlying changes in ionic homeostasis due to variations in membrane parameters in a hypoosmolar and normal media

The effects of decreasing and increasing membrane parameters by a factor 10 on the intracellular water content and ion fluxes via different pathways under the balanced state in normal medium and hypoosmolar medium of 160 mosm with NaCl reduced by 75 mM are collected in **Figure 9**. The most important conclusion here is that even rather strong variations in membrane parameters do not significantly affect the water content in the cell under the balanced state in a hypoosmolar medium. They mainly affect the dynamics of the transition. Other steady-state characteristics such as K/Na, OSOR, and U vary more significantly. It is the study of these characteristics that can help to distinguish the possible mechanisms of RVD.

**Figure 9.**
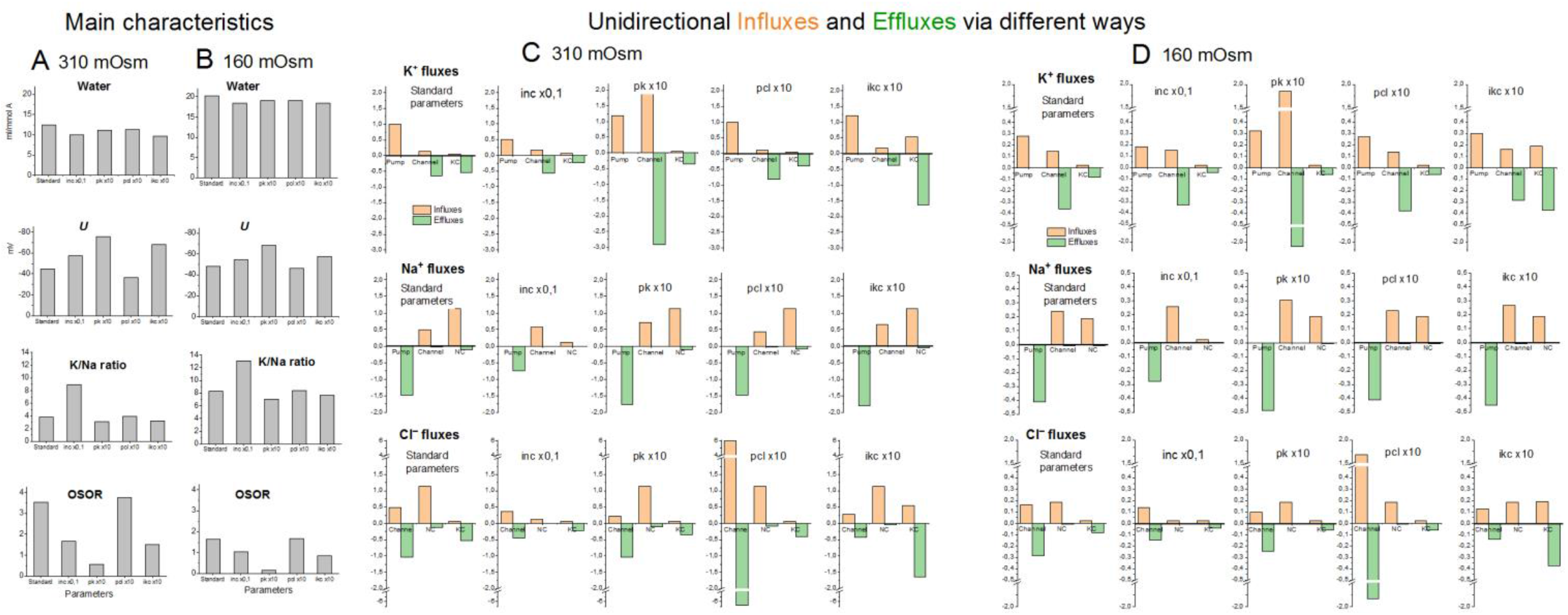
**The effects of a tenfold decreasing or increasing membrane parameters on the main characteristics of ionic homeostasis and unidirectional fluxes via different ways under the balanced state in normal 310 mOsm medium and in the hypoosmolar medium of 160 mOsm with NaCl reduced by 75 mM.** The calculation was carried out for the U937 cell model with a full set of cotransporters. The changed parameters are shown in the Figure, the others remain unchanged and are presented in Table 2, cells *B*, NC+KC+NKCC. Unidirectional flows via NKCC are not shown because they are small (see Table 4)

In accordance with the primary flux equations underlying the model, changes in partial ion fluxes are directly proportional to changes in parameters characterizing the permeability of ion channels and rate constants for ion transfer through cotransporters (*pk, pna, pcl, inc, ikc, inkcc*). Modeling show which fluxes changes more significantly and can be used for identification of the RVD mechanism using specific markers or inhibitors (**Figure 9**). Evidently, measuring K^+^ influx using Rb^+^ as its closest analog is most appropriate here.

The modeling reveals two main points. First, a change in the parameter of one ionic pathway always leads to more or fewer changes in fluxes through other pathways. This is since all channels and transporters carry ions into the common intracellular medium, and the same electrical and electrochemical gradients determine the movement of ions via channels and transporters operating in parallel. Multiple feedbacks cause multiple relationships between fluxes. Secondly, the integral characteristics of the system change much less than the parameters and fluxes in the corresponding pathways. An increase or decrease in parameters by a factor of 10 causes a decrease in cell volume only by about 20-25% both in normal and hypoosmolar media (**Figure 9, Table 5**). A further change in the parameters does not lead to a significant increase in this value. Similar variations in other parameters show that a cell cannot change its volume in a hypoosmolar medium in any range, changing the properties of channels and transporters of the cell membrane, even with a wide list of them in our model. Already here there appears a hint that there must be other ways of regulating the ionic homeostasis of cells, in addition to changing the channels and transporters of monovalent ions in the cell membrane.

**Table 5.**
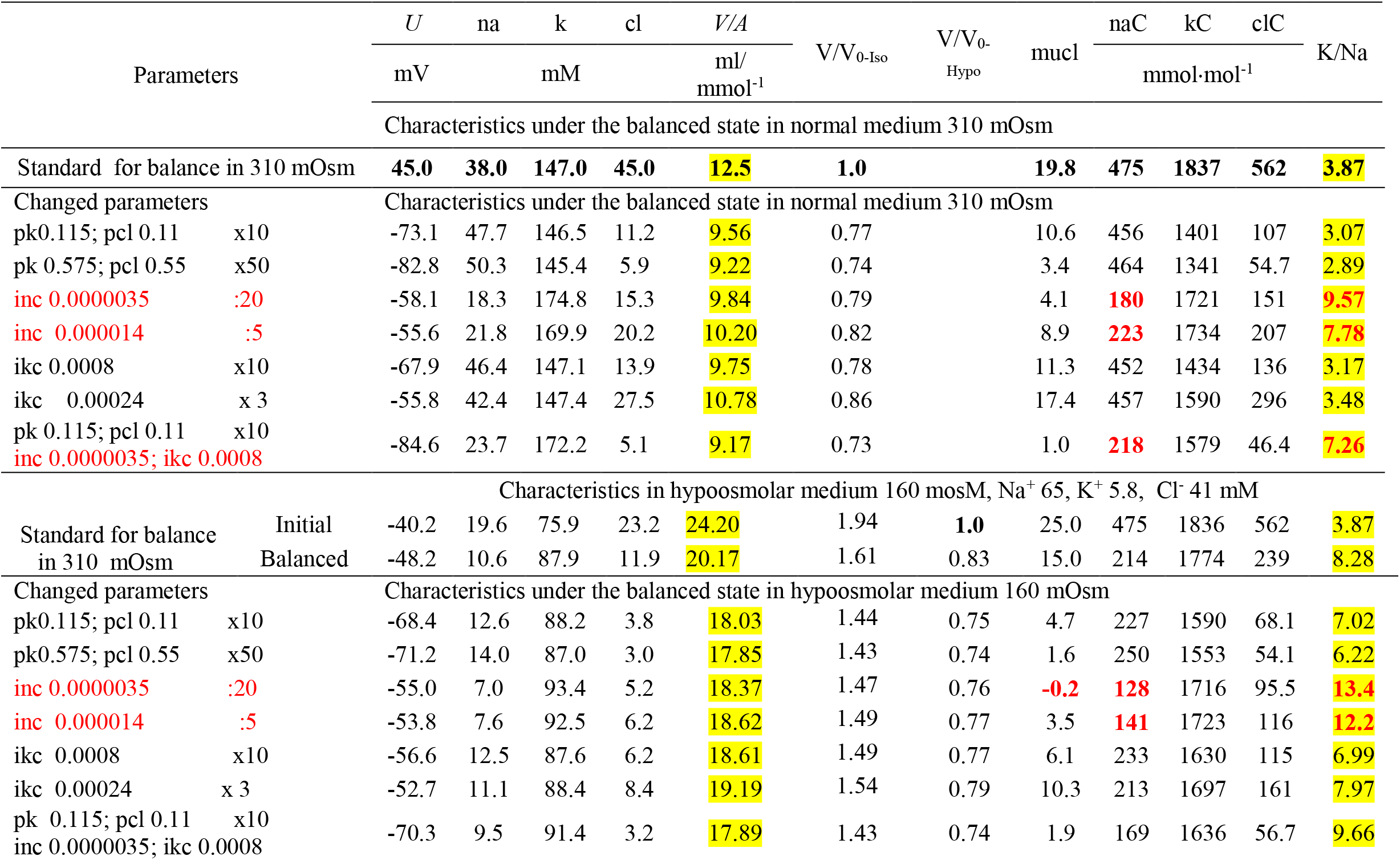
**Effect of changes in membrane parameters on the main characteristics of ionic homeostasis under the balanced state in the U937 cell model in a standard medium of 310 mOsm and in a hypoosmolar medium of 160 mOsm with a decrease in NaCl by 75 mM.** Standard parameters corresponding to the balanced state of U937 cells in the normal medium 310 mOsm, Na^+^ 140, K^+^ 5.8, Cl^-^ 116 mM are as follows: pna 0.0017; pk 0.0115; pcl 0.011; inc 0.00007; ikc 0.00008; inkcc 0.000000008; V_0-Iso_, V_0-Hypo_ – initial cell volumes in isotonic and hypotonic medium, respectively. Stronger NC effects are marked in red

#### 2.3. RVD *in living cells in the light of the U937 cell model analysis*

Our experimental data relating to U937 cells were sufficient to obtain the basic parameters corresponding to the homeostasis of monovalent ions in these cells under normal conditions. The use of the developed software and these parameters led us to unexpectedly excellent prediction of the real-time dynamics of changes in ionic homeostasis in living cells after blocking the pump. The adaptation of cells to hypoosmolar media turned out to be a more complex phenomenon in comparison with the change in ionic homeostasis after stopping the pump. **Figure 10** illustrates changes in water and ion content in living lymphoid cells of three types (K562, Jurkat, and U937) for the first 30 min and after 4h incubation of cells in hypotonic (160 mOsm) or hypertonic (510 mOsm) media which were observed in our previous study (unpublished data).

**Figure 10.**
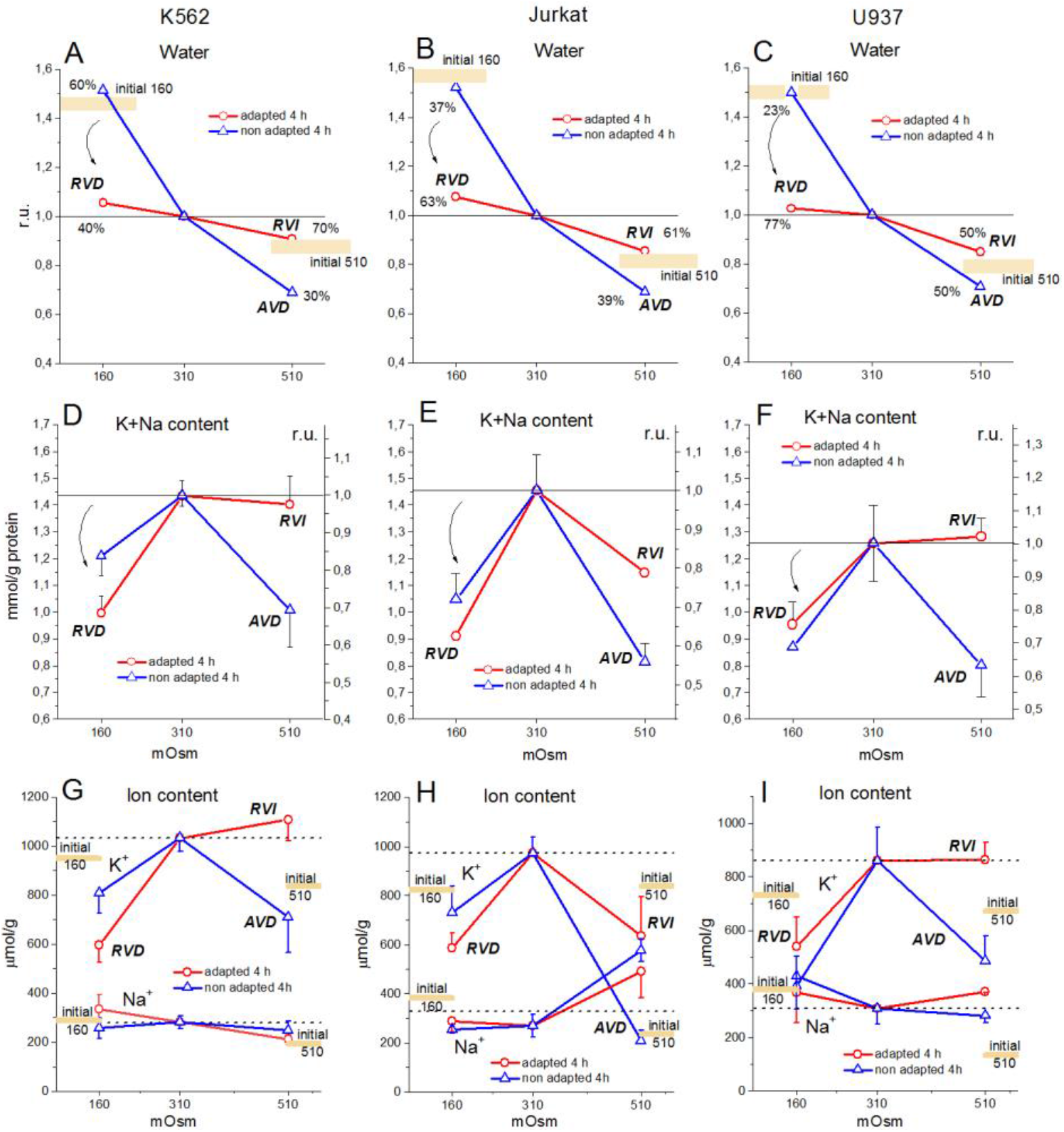
Changes in water, K^+^ and Na^+^ content in K562, Jurkat, and U937 cells at the initial 30 min and after 4 h incubation in hypo- and hypertonic medium. Extracellular osmolarity was decreased by a 75 mM NaCl reduction or increased by the addition of 200 mM sucrose to standard RPMI medium. The broad colored lines show the level of the initial values for 30 min incubation in a hypo- or hyperosmolar medium. (A-C) Water content, estimated in ml/g protein from buoyant cell density, is presented relative to cell water in standard RPMI medium.

Since cell water was assayed by measurement of the cell buoyant density, we could see that the dispersion of cells by water content when they were adapting to anisoosmolar media, both hypo- and hyperosmolar, was remarkably higher than that under standard conditions. The cell population turned out to be heterogeneous in its ability to adapt to anisoosmolar media. Therefore, the cells from upper and low parts of the band in the Percoll gradient were taken for ion assay separately. It turned out that the heavier part of the cell population consisted of 40-77 % by protein in dependence of cell species was adapted to hypotony while residual part did not (**Figure 10 A-C**). The cells adapted to the hypotonic medium contained less water and Na^+^+K^+^ than at the first moment, demonstrating RVD. The restoration of cell water in the adapted cells was nearly complete and was associated with the release of ions from the cells. However, the content of K^+^+Na^+^ did not return to the level of cells in the medium of 310 mOsm, since the external, as well as intracellular, osmolarity became 160 instead of 310 mOsm.

The relationship between changes in cell volume and the K^+^, Na^+^, and Cl‾ content have been considered already in the pioneering studies (Roti-Roti and Rothstein, 1973; Hendil and Hoffmann, 1974; Cala, 1977, Grinstein et al., 1982, 1983, Grinstein, Foskett, 1990). Since then, it has been clear that the amount of intracellular osmolytes that do not penetrate the cell membrane under normal conditions, and their charge *z*, in addition to changes in the membrane parameters, are essential player in RVD (Hoffmann et al., 2009). It has been found that the redistribution of three group of organic intracellular osmolytes is essential for RVD in many cases although their involvement in cell volume regulation is highly dependent on cell species and conditions (Kirk, 1997). Studying the pathways passing ions and intracellular organic osmolytes through the cell membrane during RVD turns out in the focus. Changes in the overall osmotic balance of a cell caused by the movement of organic osmolytes during RVD are usually not quantified. Our computation of ionic homeostasis can help to understand the possible impact of K^+^, Na^+^, and Cl‾ in RVD.

The first question is: what decrease in the content of monovalent ions should be sufficient to completely restore the water content in cells in a hypoosmolar medium? Here we need to go beyond modeling the flux balance and move on to some basic formulas. According to generally accepted concepts, two basic equations (1) and (2) determine the relationship between the content of intracellular water (*V*), the intracellular concentration of monovalent ions (Na^+^, K^+^, and Cl^-^), other intracellular osmolytes (*A*), impermeant through the plasma membrane, and their integral charge, which is always negative (Jakobsson 1980; Lew and Bookchin, 1991; Hoffmann et al., 2009 P.195-196).

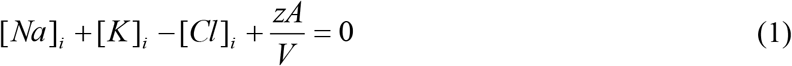

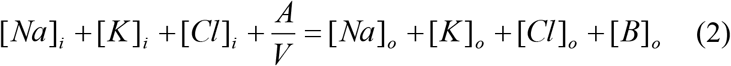

Compliance with conditions (1) and (2) means water-osmotic balance (water equilibrates much faster than ions) and much faster than ions) and integral electroneutrality of the cytoplasm. These equations underlie our calculations and are carried out not only under the balanced state in ionic homeostasis but also at any moment of the redistribution of the monovalent ions during transition from one balanced state to another irrespectively to the mechanism of ion movement across the cell membrane. *A* and *z* remain constant in our calculations. However, they can be obtained rigorously for any time point using equations (1) and (2) if the water content in cells and intracellular concentrations of Na^+^, K^+^, and Cl‾ are known for this moment.

It follows from (1) and (2) that a ratio of the volume of cells in the hypoosmolar medium to the volume of cells balanced with normal medium is determined by equation (3):

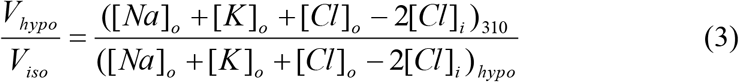

After rewriting: 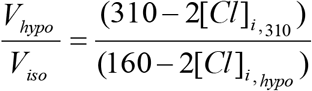

The value 2[Cl]_i,hypo_ can be found by direct measurement or by calculation for a given model. When all intracellular chloride is exhausted and [Cl]_i,hypo_ =0 the limit is achieved.

In the limit: 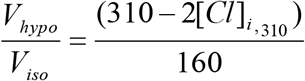

For our example of U937 cell (Table 2, cells *B*): 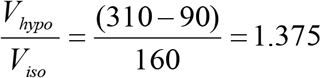

Consequently, the cell volume after RVD, caused by the maximum loss of monovalent ions at a given initial intracellular Cl‾ content, should remain 37.5% higher than in a normal medium of 310 mOsm. The volume of cells in a medium of 160 mOsm in a simple osmometer without RVD should be 1.94 times higher than normal (310/160). Thus, the swelling by 94% can be reduced due to RVD in the considered example to swelling by 37.5%.

Our experimental data for U937, K562, and Jurkat cells show that some subpopulations of these cells restore volume in the 160 mOsm medium to almost the level in the normal medium (**Figure 10**). This is an indicator that these cells can go beyond the limits described above, and use a mechanism for this, other than the changes in the membrane parameters regulating the movement of K^+^, Na^+^, and Cl‾. This can be the changes in the amount of the osmolytes considered as impermeant across the plasma membrane under normal conditions or changes in their charge. There are many direct experimental indications that the cells placed into hypoosmolar medium can loss the significant amount of organic intracellular osmolytes. These osmolytes and the pathways of their transfer across the cell membrane haven been studied intensively over the past decades. Here we show that the impact of these mechanisms in RVD can be quantified using data on changes in intracellular water, K^+^, Na^+^, and Cl‾ if obtained with appropriate accuracy.

## DISCUSSION

Despite the impressive progress in cell biology associated with advances in molecular biology, the fundamental cellular system that determines not only the water and ionic balance of animal cells, but also the electrochemical gradients of inorganic ions on the cell membrane, and the membrane potential remains in the shadows. However, this system is incredibly important for the functioning of the entire cell. The term “ionic homeostasis”, sometimes used to denote this system, seems too weak to refer to the apparatus that plays a key role in the cell physiome, that is, in the complex of physiological processes inherent in the cell which is usually not considered in areas called cellular proteome and metabolome. The electrochemical system of the cell, which includes many ion channels and carriers in the cell membrane, as well as charged intracellular osmolytes, is complex and requires computations for its analysis.

Our previous computation of the change in ionic homeostasis after stopping the pump and due to apoptosis (Yurinskaya et al., 2019) considered only the NC cotransport because of the difficulties in analyzing a system with many parameters, which must be linked to experimental data. More complex system with all main types of cotransporters NC, KC, and NKCC was computed in a previous study for U937 apoptotic cells (Yurinskaya et al., 2020). We considered how difficulties due to increasing the number of parameters can be overcome using the cotransporter inhibitors and which uncertainties remain because of inaccuracy of the primary experimental data. Here the system with all main cotransporters is used to study changes in ionic homeostasis after blocking the Na/K pump and during RVD.

In general, new calculations carried out with a wider list of cotransporters confirm that our computational approach allows us to quantitatively predict the real-time dynamics of changes in the cellular ion and water homeostasis caused by stopping the pump. Importantly, this prediction is based on the use of invariable parameters obtained for resting cells under normal conditions, without any their adjustment or fit. The accuracy of the prediction is limited mostly by the accuracy of the available experimental data.

There have been found certain differences between behavior of the model with only NC cotransporter, which is required for most cells, and the model with a several cotransporters, NC+NKCC, NC+KC, and NC+NKCC+KC. The most significant differences in the dynamics of changes in cell water content after stopping the pump for U937 cell model appear with the addition of the NKCC cotransporter. The decrease in water content in the NKCC model occurs with an extremum at a 2-hour time point, although this effect is too small to be compared with experimental data at the present accuracy of water analysis. More detailed calculations of the significance of the small amounts of impermeant osmolytes in cell environment in the reducing cell swelling after stopping the pump led us to correction of our previous views on the limited swelling of living cell under the real physiological conditions (Yurinskaya et al., 2011; Vereninov et al., 2014). The most significant result of successful prediction of the real dynamics of ion homeostasis changes after stopping the pump by calculating a model with all major cotransporters is the conclusion that the model is trustworthy.

The study of changes in ionic homeostasis caused by changes in external osmolarity is another important approach to understanding its nature and testing existing concepts and models. According to the general concept, there is some “set point” in the regulation of cell volume or cell water content. By changing the properties of channels and transporters of the plasma membrane, as well as the content of intracellular osmolytes, cells reach this set point. Our computational modeling shows that there is a “physical” RVD during the transition of cells to a hypoosmolar medium with decreasing NaCl concentration, resembling the RVD observed in living cells. This physical RVD arises with unchanged cell membrane properties due to simple changes in electrochemical ionic gradients caused by changes in the composition of the medium, rapid increase in intracellular water content, and the time-dependent changes in intracellular ionic composition. The physical RVD mask truly active regulatory processes mediated by the intracellular signaling network. Using our software allows to separate the effects of changing external osmolarity, ion composition and the properties of various kind of channels and transporters. It can be seen how the changes in the balance of the monovalent ion fluxes across the cell membrane in hypoosmolar medium may depend on the initial state of the cell.

Computation of the unidirectional fluxes as it is done in the current paper for U937 cells, allows to find the conditions when fluxes via certain species of channels or transporters monitored by ion-markers or inhibitors will be minimally masked by the fluxes via parallel pathways.

We believe that the executable file of our software is universal and can be used to calculate ion homeostasis and the balance of unidirectional flows of monovalent ions in different cells under different conditions. However, a minimal set of experimental data is required to determine the intrinsic parameters used in computation. These data include the intracellular content of cell water, Na^+^, K^+^, and Cl‾, ouabain-sensitive, and resistant components of the Rb^+^(K^+^) influx, as well as components sensitive to inhibitors of NKCC and KC cotransporters if one needs to consider cotransporters NKCC and KC. Of course, it is not easy to obtain these data with the required accuracy, especially data on the content of water and Cl‾, but a quantitative description of ionic homeostasis and balance of fluxes, as in our approach, is impossible without these data.

## CONCLUSIONS

Successful prediction of changes in ion homeostasis in real-time after stopping the pump using a model with all major cotransporters and parameters obtained for normal cells indicates the reliability of the developed computational model. The use of this model for the analysis of RVD has shown that there is a “physical” RVD associated with time-dependent changes in electrochemical ion gradients, but not with changes in channels and transporters of the plasma membrane, which should be considered in studies of truly active regulatory processes mediated by the intracellular signaling network. The developed computational model can be useful for calculating the balance of partial unidirectional fluxes of monovalent ions via all major pathways in the cell membrane of various cells under various conditions.

## Supporting information

BEZ01BC

## DATA AVAILABILITY STATEMENT

The original contributions presented in the study are included in the article/Supplementary Material, further inquiries can be directed to the corresponding author.

## AUTHOR CONTRIBUTIONS

Both authors contributed to the design of the experiments, performed the experiments, analyzed the data, and approved the final version of the manuscript and agreed to be accountable for all aspects of the work. Both persons designated as authors qualify for authorship.

## FUNDING

The research was supported by the State assignment of Russian Federation No. 0124-2019-0003 and by a grant from the Director of the Institute of Cytology of RAS.

## ACKNOWLEDGMENTS

We are grateful to Dr. Igor A. Vereninov for correcting the manuscript and suggestions for improvement.

## SUPPLEMENTARY MATERIAL

The Supplementary Material for this article can be found online at: ……………………………..

Executable file to the programme code BEZ01BC and Instruction: How to use programme code BEZ01BC.zip. This file is attached to the article electronic version.

## Conflict of Interest

The authors declare that the research was conducted in the absence of any commercial or financial relationships that could be construed as a potential conflict of interest.

## Notes

### Competing Interest Statement

The authors have declared no competing interest.

